# Multiobjective design of growth media with genome-scale metabolic models and Bayesian optimization

**DOI:** 10.64898/2025.12.12.693982

**Authors:** Nicola Hallmann, Catalina Guerra-Cornejo, Karl Burgess, Charlotte Merzbacher, Diego A. Oyarzún

## Abstract

The optimization of culture media is critical for improving the efficiency and cost of cellular production systems. Traditional approaches often rely on extensive experimental trials or statistical methods, which can be both costly and time-consuming. Here, we present gsMOBO, a computational approach for media design that integrates genome-scale metabolic models with multiobjective optimization. Our method employs Flux Balance Analysis coupled with a top-layer Bayesian optimization routine for efficient exploration and optimization of nutrient combinations across high-dimensional spaces. We show that our method finds optimal medium formulations along a Pareto front balancing growth, production and cost of medium components. We illustrate the approach in models of *Escherichia coli* engineered to produce antibody fragments, as well as *Bacillus subtilis* strains that synthesize cyclic lipopeptides. Our results show that gsMOBO identifies media compositions and Pareto-optimal trade-offs consistent with prior experimental work. Our method provides a broadly applicable tool for the design of cost-effective and productive culture media, and offers a route to accelerate medium development in biomanufacturing.

## 1 Introduction

The performance of production strains is heavily influenced by the composition of the culture media, which must align with the metabolic demands of the host to maximize growth and productivity. Optimal media formulations are also critical for managing costs and determining the economic viability of a bioprocess [1, 2]. Media cost is a significant barrier for scale-up production of small molecules [3] and protein products [4, 5], where media components tend to dominate the upstream costs [6].

Traditional approaches to media design combine host-specific knowledge with trial-and-error experimentation and statistical design of experiments [7, 8]; these techniques are resource-intensive and become infeasible as the number of medium components increases. As a result, many studies have employed several computational tools for media optimization [8, 9, 10, 11]. The use of genome-scale metabolic models (GEM) has been particularly successful for optimizing production in a range of microbial [12, 13] and mammalian hosts [14, 15, 16]. Despite the broad adoption of GEMs [17], their successful use in media design often relies on case-specific analyses that are difficult to generalize across different hosts and products [18, 19]. A recent study proposed an algorithm to design nutrient supplementation strategies [20] and a growing body of work has focused on integrating GEMs with machine learning algorithms for media design [21, 22, 23], thanks to the improved predictivity observed in other metabolic engineering tasks [24, 25, 26, 27].

Here, we present gsMOBO as a general strategy for media design based on genome-scale metabolic modelling in tandem with multiobjective Bayesian optimization. The approach allows computing Pareto-optimal media compositions that trade-off growth rate, production flux, and the cost of medium components. Our strategy is based on a standard Flux Balance Analysis (FBA) solver wrapped into a top-layer optimizer that navigates a high-dimensional space of media components. To avoid the computational pitfalls of gradient-based optimizers, which require computing FBA solutions at many locations of the input space, we employed Bayesian optimization [28] for efficient sampling and improved speed. Bayesian optimization is well suited for problems with objective functions that are expensive to compute [29], and has found applications in a range of synthetic biology tasks such as protein engineering [30], gene circuit design [31] and active learning for strain design [32, 33].

Our results show that gsMOBO can robustly find two- and three-dimensional Pareto fronts by searching over up to 13 media components simultaneously. We illustrate the utility of the approach for maximization of growth rate and production with minimal media cost, using GEM models for production of antibody fragments in *Escherichia coli* [18] and synthesis of lipopeptides in *Bacillus subtilis* [34]. The implementation of gsMOBO has been designed to interface seamlessly with current GEM model specifications and the COBRApy package [35]. Our work presents a widely applicable framework for computational screening of media compositions with applications in a breadth of metabolic engineering campaigns.

## 2 Results

### 2.1 Multiobjective Bayesian optimization of genome-scale metabolic models

We consider the optimization of medium composition for strains described by a genome-scale metabolic model (GEM) using the following relations:

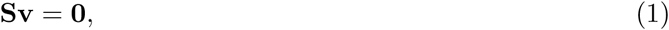

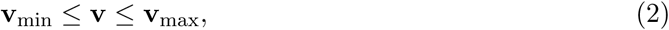

where **S** is the stoichiometry matrix of a metabolic network with *m* metabolites and *n* reaction fluxes comprised in the vector **v** [17]. Each reaction flux is constrained by physiological or thermodynamic bounds **v**_min_ and **v**_max_. To formulate media design as an optimization problem, we split the constraint (2) into two sets of component-wise inequalities:

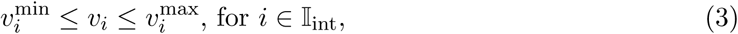

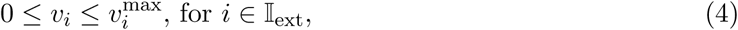

where *v_i_* is the *i*-th reaction flux and the index sets 𝕀_int_ and 𝕀_ext_ correspond to the internal reactions and nutrient import reactions, respectively. The bounds on the internal fluxes in 𝕀_int_ can be estimated from various experimental data [36]. The upper bounds on nutrient import reactions in 𝕀_ext_, on the other hand, are employed to model the growth media. If a nutrient is absent from the growth media, its corresponding flux bound is set to 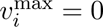.

We define the media components to be optimized (i.e. the decision variables) via the index set 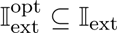, and formulate the media design task as the following multiobjective optimization problem:

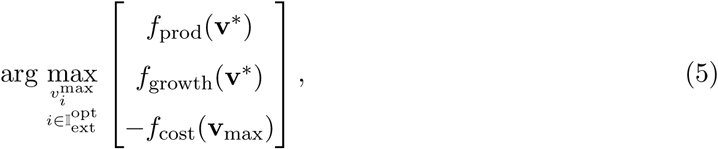

where *f*_prod_, *f*_growth_, and *f*_cost_ are objective functions that quantify production performance, strain growth rate, and cost of medium components, respectively. The cost objective (*f*_cost_) is computed from the components supplied in the medium (**v**_max_) linearly weighted by the price per component. The production and growth objectives are computed as linear objectives depending on the flux vector **v**^∗^, which is a solution of a standard Flux Balance Analysis (FBA) problem [17]:

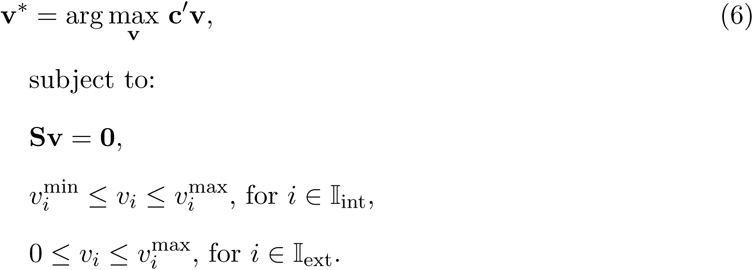

In the FBA problem, the vector **c** contains weights to model a combination of the rate of biomass synthesis alongside the production flux in specific pathways of interest. The approach in Eq. (5) is a bi-level optimization problem, whereby an outer optimizer searches for combinations of exchange flux bounds that maximize two or more objective functions, which in turn are computed via an inner FBA optimizer; from a geometric standpoint, this strategy can be seen as optimizing the boundary facets of the convex polyhedron defined by the GEM.

As illustrated in Figure 1A, solving the multiobjective optimization problem requires computing FBA solutions at many points of the media space. But as the number of components grows, the search space grows combinatorially large and the solution becomes computationally infeasible with conventional optimization approaches. To resolve this, we employed Bayesian optimization [29], a global optimization technique for objective functions that are expensive to compute. It models the objective function using a probabilistic surrogate, typically a Gaussian Process (GP), which provides both mean predictions and uncertainty estimates. An acquisition function then guides the selection of the next evaluation point by balancing exploration and exploitation, so as to sample uncertain regions and refine predictions near promising areas of the media component space. After each iteration, the GP is updated with the new evaluation data, refining the posterior distribution over the objective. This iterative process can efficiently converge toward the global optimum with a reduced number of function evaluations.

**Figure 1:**
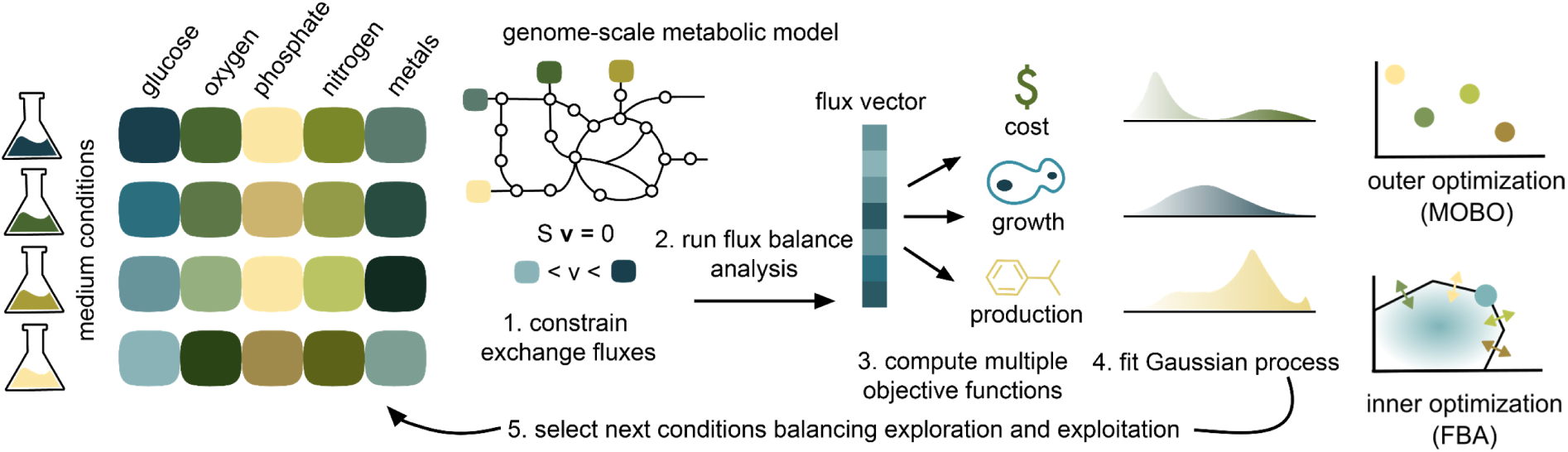
Optimization of medium components with gsMOBO. Schematic of the gsMOBO framework. Medium compositions of various concentrations can result in different growth and production phenotypes. Exchange fluxes in a genome-scale metabolic model of the host are constrained based on the medium conditions and flux balance analysis (FBA) produces an optimal flux vector. This is used to compute the growth and production objectives for the Bayesian optimization loop; the medium cost is computed from the upper bounds of media components. The Bayesian optimization iteratively fits a Gaussian process model of the medium condition space and uses an acquisition function to select the next medium conditions to sample, balancing exploration of the large design space with exploitation towards a global optimum. The outer optimization loop selects points in the medium condition spaces (colored dots); each of these points corresponds with a change in the bounds of the high-dimensional flux cone (colored arrows), which affects the optimal FBA solution found by the inner optimization (blue dot).

gsMOBO outputs a set of nutrient import fluxes that optimally balance the different objectives, from where designers can choose a preferred media based on the trade-off between production, growth and cost. These nutrient combinations lie along a Pareto front, which describes the best possible trade-offs between conflicting goals in the sense that no objective can be improved without making another one worse.

### 2.2 Optimization of growth rate and media cost in *Escherichia coli*

We first tested gsMOBO for the design of culture media that balance growth rate against the cost of media components, using the iML1515 model for wild type *Escherichia coli* [37] as a test case.

To this end, we formulated the problem as joint maximization of two objective functions *f*_growth_ = **c**^′^**v**^∗^ and *f*_cost_ = −**z**^′^**v**_max_, where **c** is the vector of weights informed by biomass composition, and **z** is a price vector for each medium component sourced from commercial vendors (Supplementary Table S1). We employed the iML1515 model with an M9 media supplemented with essential trace metals; omission of any of these metals resulted in zero predicted growth rate in iML1515. We employed gsMOBO to optimize ten media components: ammonium (NH_4_^+^), calcium (Ca^2+^), chloride (Cl^−^), glucose, magnesium (Mg^2+^), potassium (K^+^), phosphate (PO_4_^3−^), sodium (Na^+^), sulfate (SO_4_^2−^), and oxygen; these decision variables include all components of the standard M9 medium, except for the trace metals.

The method successfully converged to media combinations that trade off growth rate and medium cost (Figure 2A). The two objectives form a near convex Pareto front and we observed that medium compositions sampled later in the algorithm tended to perform closer to the Pareto-optimal solutions. As gsMOBO progressed through its iterations, we observed an improvement in the best growth/cost ratio (Figure 2B). Despite converging to the Pareto front, the algorithm was able to continue sampling broad regions of the objective function space across all iterations. This reflects the key advantage of Bayesian optimization as compared to gradient-based methods that tend to get trapped in local optima.

**Figure 2:**
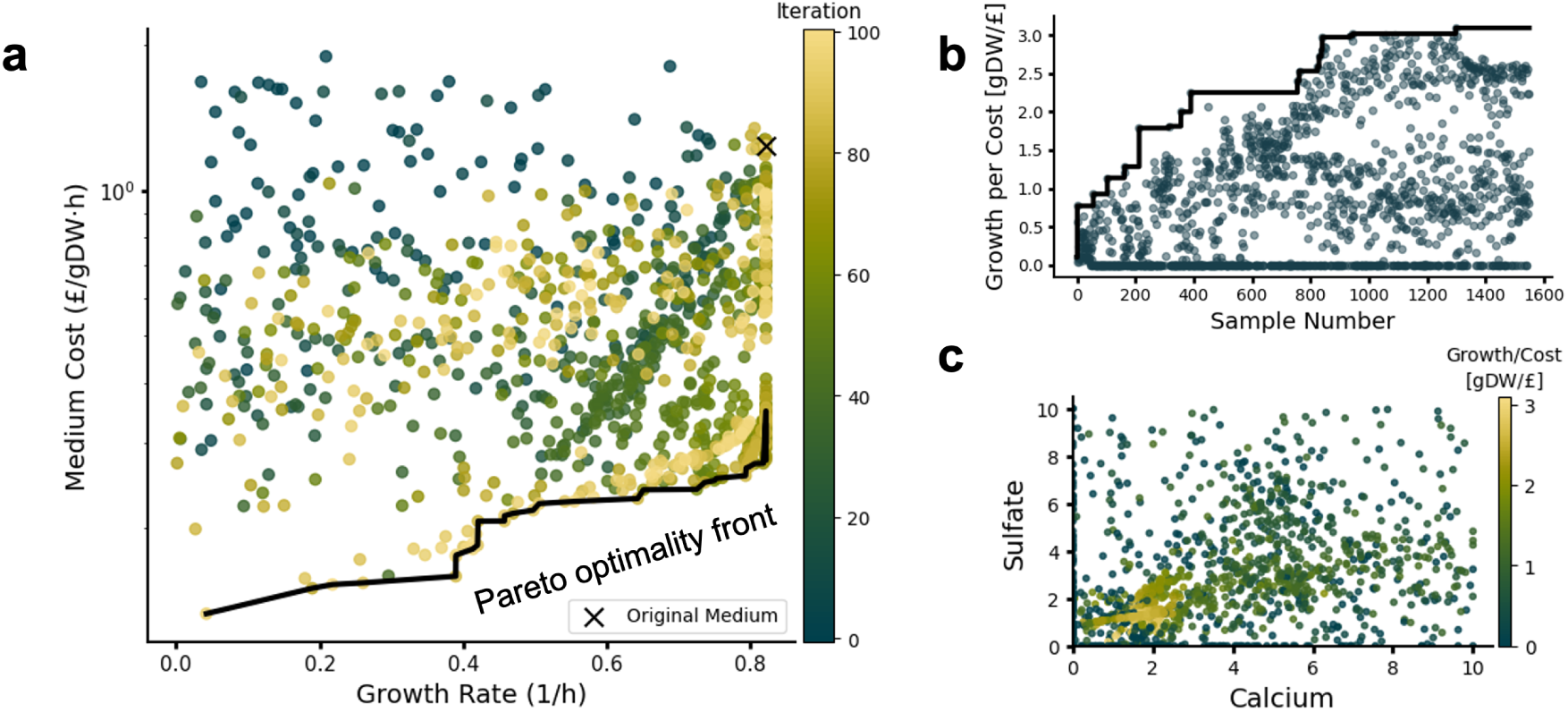
Optimization of wild type *Escherichia coli* medium. (**A**) Pareto front between optimal growth rate and medium cost computed for the *E. coli* iML1515 model using ten medium components as decision variables; the color bar denotes the iteration number in the gsMOBO run. gsMOBO samples get closer to the Pareto front as it progresses through its iterations. (**B**) Exemplar run quantified by the growth/cost ratio. The black line is the best growth per cost sample seen so far across the run. The algorithm continues to sample across the space throughout the run. (**C**) Calcium and sulfate, two medium components, and their relationship to growth and cost. The samples of gsMOBO converge on a relatively small region of the space with high growth per cost, though widespread sampling continues throughout the run. The algorithm was initialized with 50 random medium compositions and ran for 100 iterations, sampling 15 medium compositions in each loop.

The results in Figure 2A indicate that samples are enriched for media with good growth rates above 0.6 h^−1^, with ∼ 8% of sampled media formulations leading to the maximum growth rate of 0.822 h^−1^. The cheapest of these maximal growth formulations costs 22.3% of the initial M9 formulation, which suggests that cost savings are feasible without sacrificing the growth rate. To verify if gsMOBO sampled the entire input space while also more densely exploring regions close to the objective optimum, we visualized two-dimensional cross sections of the full media component space (Figure 2C). This indicates that the optimizer effectively samples data points spread across the whole possible input space but concentrated in the regions of the space with high growth/cost ratio.

### 2.3 Triple objective optimization of antibody production in *Escherichia coli*

To test gsMOBO in a more challenging task, we focused on optimization of recombinant antibody production in *E. coli*, whereby media costs can be a barrier for commercial scale-up [6]. We employed a literature model of *E. coli* engineered to produce anti-EpCAM extracellular domain single-chain variable antibody fragments [18]. This work employed GEM analysis to identify amino acid supplementation for improved growth and production. The model was built on top of an earlier GEM for *E. coli* iJO1366 with a reaction producing the plasmid containing the antibody gene (Figure 3A), using an FBA objective function that maximizes a combination of biomass and product synthesis in a 99:1 ratio (details in Methods).

**Figure 3:**
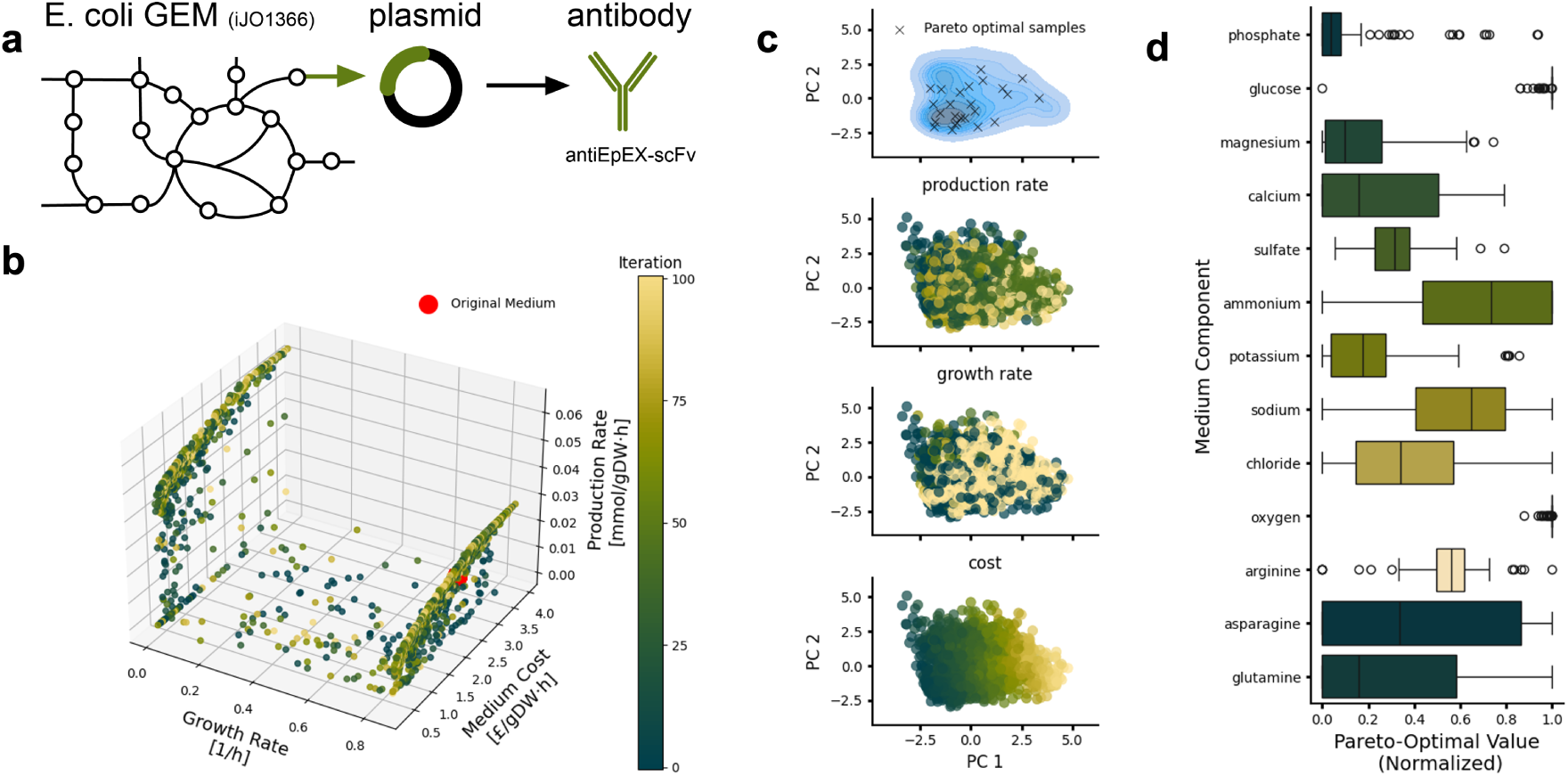
Triple-objective optimization of antibody production in *Escherichia coli*. (**A**) An *E. coli* GEM (iJO1366) modified with a reaction to produce a plasmid containing the antibody antiEpEX-scFV was sourced from the literature [18]. (**B**) Three-dimensional Pareto front (production rate, growth rate, medium cost) computed with gsMOBO using 13 components as decision variables. (**C**) Principal component analysis (PCA) of medium compositions, labeled by their Pareto optimality, production rate, growth rate, and medium cost. PCA plots were computed on the 13-dimensional vectors of media components for all gsMOBO samples. The top panel shows the Pareto-optimal compositions overlaid with a density estimate of all other compositions. The other panels show the individual gsMOBO samples. (**D**) Box plots of all Pareto-optimal normalized medium compositions. Some components (glucose, oxygen) are optimized to a small range, while other components (asparagine, glutamine) can take large ranges while remaining Pareto-optimal. The algorithm was initialized with 50 random medium compositions and ran for 100 iterations, sampling 15 medium compositions in each loop.

In addition to biomass synthesis (*f*_growth_) and media cost (*f*_cost_), we considered a third objective representing production of the antibody fragment *f*_prod_ = **p**^′^**v**^∗^, where **p** contains suitable weights to model the different precursors requires to assemble the product. As decision variables, we considered the same ten media components employed in the iML1515 model in the previous section, plus the amino acids glutamine, arginine and asparagine. Supplementation of these amino acids has been shown to support antibody fragment production [18]. We first optimized all three pairwise combinations of objective functions (Supplementary Figures S1-S3). In all cases, gsMOBO successfully converged to solutions predicted to have a better trade-off than predicted for the baseline media.

We next performed a triple objective optimization of growth, antibody production and media cost. The algorithm was able to effectively identify a 3-dimensional Pareto front (Figure 3B). It effectively sampled broad regions of the objective space, yet we observed a concentration of samples along two regions associated with near maximal growth and low production, and minimal growth and high production. Within these two regimes, gsMOBO sampled the entire range of feasible production rates, but only few mid-growth rate compositions were found and most of them led to low production. The same pattern was observed for the double optimization of growth rate and production rate (Supplementary Figure S1). This is likely a consequence of the FBA optimization, which inherently favors phenotypes at the boundary of the flux cone due to its linear programming formulation. Better coverage of the mid growth region can likely be achieved with an increased number of samples or calibration of the acquisition function that balances exploration and exploitation in the Bayesian optimizer.

The Pareto front Figure 3B captures pronounced trade-offs between the different design objectives. For example, the best production-cost trade-off was achieved at ∼69.4% of the maximal production rate, with medium costs that are only ∼15.3% of those of the medium that supports the maximal production rate. This illustrates that fully optimizing one objective might incur disproportionate costs on another one. The results in Figure 3B also predict 0.0275 mmol gDW^−1^h^−1^ as the maximum production rate at near-maximal growth. This is a 31% improvement with respect to the prediction with M9 medium supplemented with amino acids identified in the original work by Behravan et al. [18] at a 22.2% higher cost. To explore this result further, we ran the triple objective optimization with only the three supplemented amino acids as decision variables, leaving all other components fixed to the M9 baseline. The maximum production rate identified in this scenario was 0.0272 mmol gDW^−1^h^−1^ at 36.8% higher cost than M9, which is comparable to the optimal solution at maximum growth previously found. Notably, in this case, gsMOBO did not identify the low growth, high production regime as part of the Pareto front, which supports previous experimental findings [18] and suggests that amino acid supplementation ameliorates the performance trade-offs.

To better understand the structure of the sampled solution space and to visualize where the Pareto-optimal media lie in relation to the broader set of samples, we performed Principal Component Analysis (PCA) for the media compositions (Figure 3C). The PCA projection reveals that the Pareto-optimal samples occupy a well-defined subregion of the design space, indicating convergence of the gsMOBO toward specific compositional profiles. Production rate, growth rate, and medium cost exhibit distinct gradients across the principal components, suggesting that the underlying variation in these objectives is well captured by the first two principal components.

To understand how specific components contribute to the optimized trade-offs, we analyzed the distribution of medium components across Pareto-optimal samples (Figure 3D). Several components, such as calcium, chloride, and ammonium, show broad distributions, suggesting that optimal performance can be achieved across a range of concentrations for these nutrients. In contrast, components like phosphate and glucose show tight distributions near the upper bound, reflecting their known role in achieving high growth and production. Interestingly, amino acids such as asparagine, glutamine, and arginine exhibit more variable usage patterns, possibly pointing to flexibility in nitrogen source selection across different trade-offs. These results highlight how Bayesian optimization can uncover not only optimal combinations but also the sensitivity and robustness of each medium component within multi-objective constraints.

### 2.4 Optimization of surfactin production in *Bacillus subtilis*

As an additional test of gsMOBO in more complex metabolic engineering campaigns, we applied it to the production of surfactin in *Bacillus subtilis*. Surfactin is a cyclic lipopeptide naturally produced in trace amounts by *B. subtilis* (Figure 4A), with multiple applications as antimicrobial, antiviral, and emulsifying agent [38]. Optimizing growth media for surfactin production is particularly challenging because its biosynthesis is carried out by a non-ribosomal peptide synthetase with multiple precursor pathways from branched-chain fatty acids and amino acids. Owing to the high production costs and low yields [39], various studies have explored media design as a strategy to improve titer [34, 40, 41, 42].

**Figure 4:**
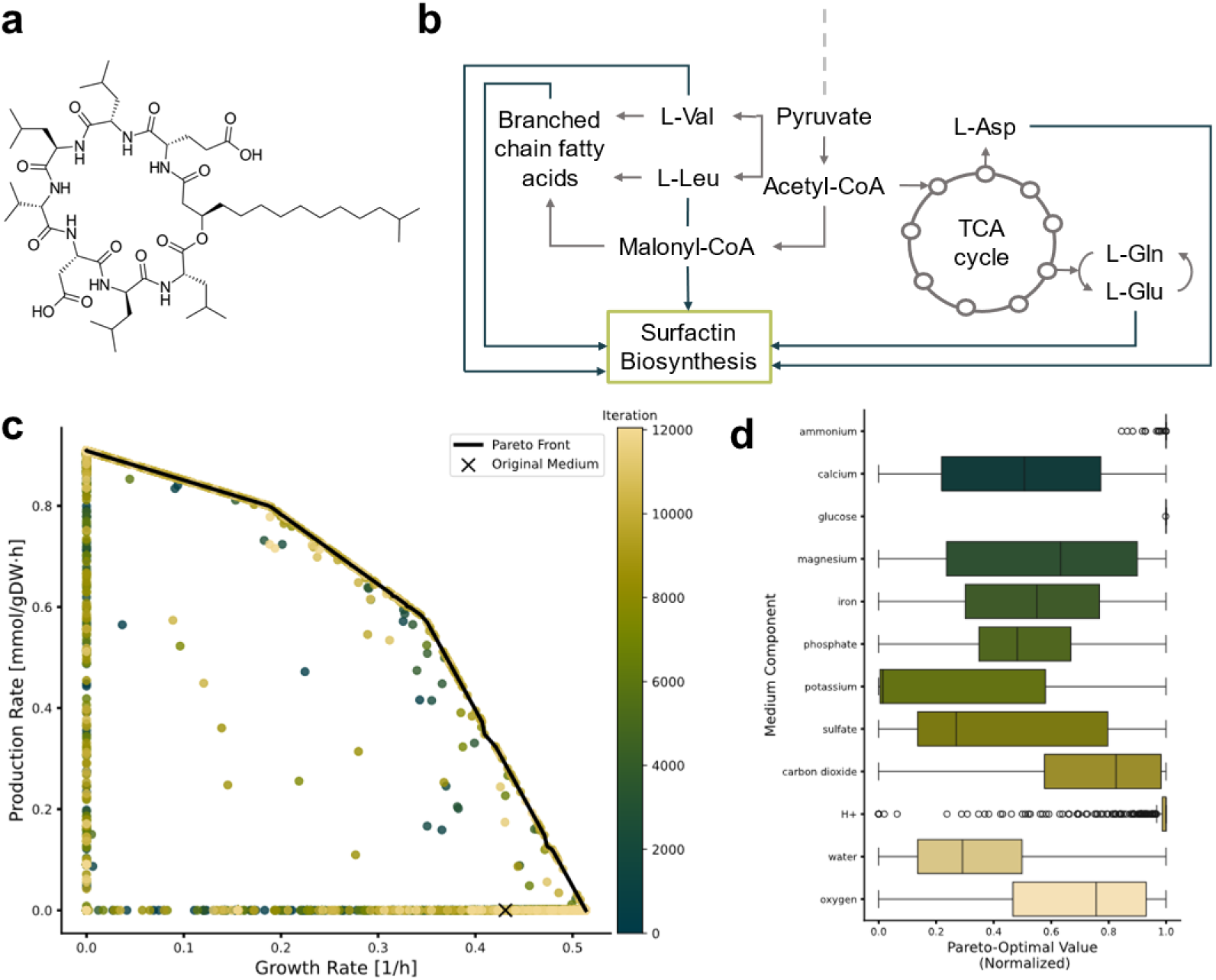
Double-objective optimization of surfactin production in *Bacillus subtilis*. (**A**) Structure of surfactin D (Equation 9) showing its hydrophobic fatty acid tail and hydrophilic amino acid ring (L-Glu, L-Leu, D-Leu, L-Val, L-Asp, D-Leu, L-Leu). (**B**) Precursor pathways present in the iBsu1103 GEM involved in surfactin production. (**C**) Pareto front between growth rate and surfactin D production rate obtained with gsMOBO with M9 medium components as decision variables. To increase coverage of the front, we ran gsMOBO six times with *N* =200 iterations per run; the first run was initialized with 50 random compositions and the next runs were initialized using the Pareto-optimal samples from the previous run. (**D**) Box plot of normalized media compositions in the Pareto front.

We employed gsMOBO to perform a double-objective optimization of growth and surfactin production. To this end, we extended the iBsu1103 GEM for *B. subtilis* [43] with a sink reaction that links the different pathways involved in surfactin assembly (Figure 4B). As objective functions for gsMOBO, we employed the default biomass reaction (*f*_growth_) included in the iBsu1103 model and production flux of surfactin D (*f*_prod_). We ran gsMOBO using 13 media components as decision variables: ammonium (NH_4_^+^), calcium (Ca^2+^), glucose, magnesium (Mg^2+^), phosphate (PO_4_^3−^), potassium (K^+^), sulfate (SO_4_^2−^), iron (Fe^3+^), oxygen, protons (H^+^), water and carbon dioxide (CO_2_); these include most M9 media components (except chloride and sodium), iron as a essential trace metal to the iBsu1103 model, and oxygen provided by the stirring conditions. Chloride and sodium were excluded as they do not appear in the iBSu1103 biomass reaction or the surfactin production reaction. Protons, carbon dioxide and water were also considered as decision variables due to their presence in the fatty acids elongation cycle and surfactin reactions. The results in Figure 4C show that gsMOBO was able to identify a Pareto front between growth and surfactin production, which qualitatively resembles experimentally determined trade-offs between *B. subtilis* growth and surfactin titer [34].

Unlike our previous case studies, where a single gsMOBO run was sufficient to find the Pareto front, in this case, we needed to run the algorithm recursively to get enough coverage of the front. We ran gsMOBO six times in total, initializing the first run with randomized data and using the previous Pareto optimal sample as the starting point for the next run. Even with this recursive strategy, only ∼3.6% of samples in Figure 4C were found in the Pareto front, and nearly 94.8% of samples were located at the null growth or minimal production boundaries. This difficulty in finding Pareto-optimal media compositions likely results from the dense connectivity of the various metabolic pathways required to synthesize surfactin. Nevertheless, gsMOBO was able to find medium compositions that reach up to 25% higher growth rates, compared to the original M9 medium. These media combinations not only are able to reach higher growth, but also produce surfactin rates above the minimal boundary. This case study shows that it’s feasible to reach higher titers of surfactin by only modifying certain components of a minimal mineral medium.

The distribution of media components along the Pareto-optimal compositions (Figure 4D) shows that most components display wide variation, without an obvious enrichment pattern along the growth-production trade-off. However, glucose and ammonium display much narrower distributions across the Pareto front; in particular, glucose consumption is predicted to be at maximum for all Pareto-optimal compositions. This may be explained by the presence of carbon and nitrogen in both the biomass objective and the surfactin production flux through L-Val, L-Leu, L-Asp and L-Glu, which pushes their consumption to the upper limit. Other components that appear only in the biomass reaction and are absent from the surfactin reaction display much wider distributions, possibly because they are involved in multiple precursor pathways such as amino acid synthesis and fatty acid elongation cycle. These results highlight the utility of gsMOBO to explore a wider range of supplementation strategies, for example, to systematically explore the benefits of ammonium against combinations of key amino acids as nitrogen source [41].

## 3 Discussion

In this work, we introduced gsMOBO as a general method for optimizing culture media compositions *in silico*. By iteratively refining a nonparametric model based on FBA predictions, our approach can identify medium formulations that balance growth, production flux, and component cost. In our test cases in *E. coli* and *B. subtilis*, gsMOBO was able to uncover Pareto-optimal trade-offs between these competing objectives. Traditional approaches to media design often rely on Design-of-Experiments approaches or nutrient supplementation [3, 8, 44]. Unlike these empirical strategies which are typically limited to a small number of components, gsMOBO can explore high-dimensional design spaces and uncovers synergistic effects between medium components that might otherwise be missed.

We found that gsMOBO consistently revealed biologically meaningful trade-offs, such as the inverse relationship between antibody production and cellular growth rate. This observation echoes established metabolic principles and supports earlier findings that optimizing for one performance metric often entails sacrificing another [18, 45]. Moreover, our results highlight the generalizability of this approach across metabolic models and use cases. By applying gsMOBO to two distinct GEMs and objective functions, we show that this framework is model-agnostic and adaptable, aligning with growing calls for more flexible, data-efficient approaches to medium design [8, 36]. As a multi-objective optimizer, gsMOBO treats production, growth and cost as independent targets, which contrasts with other approaches that rely on scalarized objective functions [20]. The multi-dimensional optimality of metabolism has been thoroughly explored in natural systems [46]. In the case of metabolic engineering campaigns, multi-objective optimization has been employed to extend traditional FBA optimization to competing targets [47], as well as for the design of gene knock-out and knock-in strategies [48]. gsMOBO provides a novel application of multi-objective optimization of GEMs, focused on the design of medium components.

Our approach relies heavily on the accuracy and completeness of the underlying GEMs. While models such as iML1515 and iJO1366 are well-validated [37, 49], they still simplify cellular metabolism and omit regulatory or kinetic constraints. Further extensions could include the use of dynamic FBA to model various bioreactor operating modes, and the inclusion of flux variability analysis (FVA) or flux sampling to improve performance by providing flux distributions for each medium composition [24, 50]. Furthermore, although Bayesian optimization is sample-efficient, it is not guaranteed to find the global optimum and, as we observed in the case of surfactin production, its performance may degrade with production tasks that depend on many precursor pathways. The integration of gsMOBO with experimental workflows is an exciting avenue for future work. Some previous work has integrated multiple experimental information sources of varying quality to rapidly optimize medium composition [51]; the same approach could be applied if there are multiple GEMs or a mixture of GEM simulations and experimental data. Automated, high-throughput fermentation systems could serve as physical validation platforms, enabling closed-loop optimization where experimental data refines the surrogate model in real time.

In conclusion, our study demonstrates that gsMOBO provides a general framework for the multi-objective optimization of culture media. By design, the Bayesian approach balances exploration of uncharted regions with exploitation of known high-performing areas of the media component space [52, 53]. This allows for a systematic screening of the multi-dimensional space of medium components, revealing trade-offs that are key for effective bioprocess engineering and precision fermentation.

## 4 Methods

### 4.1 Bayesian optimization

Our implementation of gsMOBO is built on top of the multi-objective Bayesian optimization tools available in BoTorch (version 0.16.0) in Python (version 3.12.6), using the qParEGO algorithm [54] and qLogExpectedImprovement acquisition function with an augmented Chebyshev scalarization. As input, gsMOBO receives a COBRApy genome-scale metabolic model (GEM) together with:

- data structures defining the decision variables (medium components), the upper and lower bounds for the max allowed influx, and associated costs.
- the objectives for the outer optimization loop; and
- the FBA objective function used in the inner loop

gsMOBO allows users to specify the number of randomly sampled media compositions used for initialization, the number of iterations for gsMOBO, and the batch size (number of medium compositions sampled per iteration).

The optimizer is initialized by evaluating a randomly sampled set of media compositions with respect to the chosen objectives. At each gsMOBO iteration, the surrogate model is initialized and trained using all, 0-1-normalized, previously evaluated media compositions and their corresponding objective values. Production and cost objectives were 0-1 normalized, and cost was inverted by computing (1 – normalized cost) so that all objectives are optimized in a consistent orientation; the growth objective was left unnormalized.

The multi-objective problem is scalarized using an augmented Chebyshev scalarization, and the scalarized objective is evaluated on the surrogate posterior. A batch of candidate media compositions is obtained by defining a qLogExpectedImprovement acquisition function for each and optimizing the entire list over the 0–1 design space. These candidates are then denormalized, converted into COBRApy-compatible dictionaries, and evaluated via FBA to obtain the flux vector **v**^∗^ needed to evaluate the two objectives (*f*_growth_ and *f*_prod_) as well as via calculating the medium cost. Each newly evaluated medium composition is appended to the dataset and used in the next iteration. After completion of all iterations, the Pareto-optimal media compositions are identified.

The gsMOBO results for *E. coli* models were obtained on a Lenovo ThinkPad T14 Gen6 with an AMD Ryzen AI 7 PRO 350 processor (8 cores, 16 threads; base clock 2.0 GHz). Runtimes for *E. coli* were 56 min and 92 min for the double and triple objective optimizations in Figure 2-3, and 78 min, 64 min, 94 min and 104 min for the double and triple objective optimizations in Supplementary Figures S1–S4. Runtime for each run of the *B. subtilis* double optimization in Figure 4 were 218 min, 279 min, 347 min, 360 min, 394 min and 399 min, respectively, using an HP EliteBook 840 (10 cores, 12 threads; base clock 1.60 GHz). Runtime improvements can be further made with GPU acceleration included in gsMOBO.

### 4.2 Genome-scale metabolic models

#### 4.2.1 Wild type *Escherichia coli*

As a proof of concept, we employed the iML1515 model for *E. coli* [37], with its default medium replaced with minimal M9 medium for baseline comparisons. We considered all M9 media components (ammonium, calcium, chloride, glucose, magnesium, phosphate, potassium, sodium and sulfate) and oxygen as decision variables, and fixed the upper bounds for the remaining components in M9. Specifically, we fixed the uptake rates for the trace metals manganese, iron, zinc, nickel, copper, cobalt, and molybdate at a low value sufficient not to impair predicted growth. This approach allowed us to model these components as being in excess and let the FBA optimization determine the uptake rate. We employed the default biomass objective function included in iML1515 for the FBA optimization.

In the outer Bayesian optimization loop, we employed box constraints on the maximum uptake fluxes that were considered as decision variables by setting 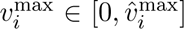 for 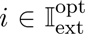 in the notation of Eq. (5). The values for 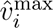 were set above those for M9, all of which can be found in Supplementary Table S1. The upper bound of oxygen uptake was set to 20 mmol gDW^−1^h^−1^ [55], placing it at a mid-point of the observed oxygen uptake rate range by various *E. coli* strains.

#### 4.2.2 Production of antibody fragments in *Escherichia coli*

We employed a GEM developed by Behravan et al [18] that was based on the iJO1366 model for *E. coli* [56] extended with plasmid-based expression antiEpEX-scFv (anti EpCAM extracellular domain single-chain variable fragment). As a reference for the optimization, we employed the same M9 growth medium as in the wild type *E. coli* model, supplemented with the amino acids that Behravan et al found to optimally support growth and protein production (9.9 mM of glutamine, 9.5 mM of arginine and 6.1 mM of asparagine). The bounds for amino acid supplementation can be found in Supplementary Table S1. The FBA objective function was defined as the weighted average of the biomass reaction and the production flux in a 99:1 ratio.

#### 4.2.3 Production of lipopeptides in *Bacillus subtilis*

We employed the iBsu1103 GEM for *Bacillus subtilis* developed by Henry et al [43], extended with reactions for four types of surfactin:

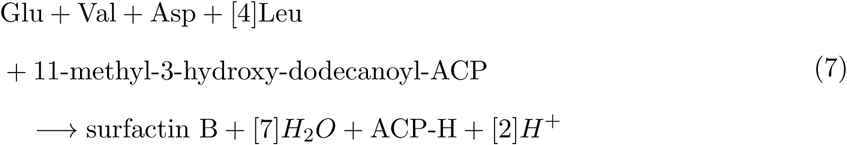

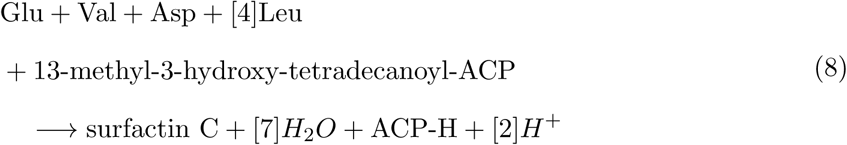

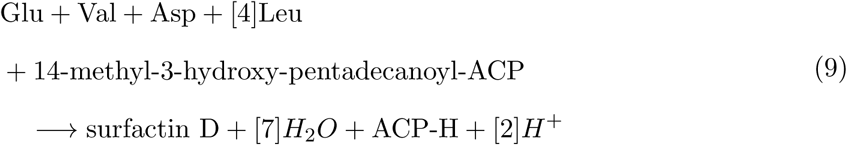

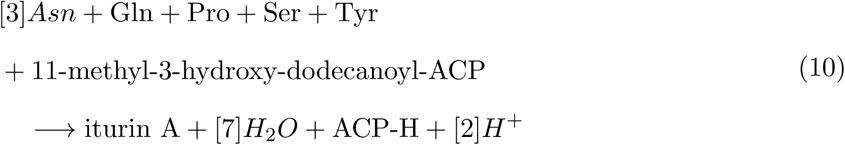

These reactions model the supply of amino acids and the fatty acid tail required for surfactin assembly. Regarding the amino acid ring, each amino bond produces a molecule of water. As for the fatty acid tail, its length can vary between 12 and 17 carbons [57], we employed tails from PubChem data for simplicity. In iBsu1103 model, the above-mentioned fatty acid tails are only available with an acyl carrier protein (ACP) linked as part of the elongation cycle. We used these variations by generating the ACP group as a byproduct. In summary, we considered the corresponding amino acids and the fatty acid-ACP complex as reactants and each surfactin type, water, 2 hydrogen ions and an ACP group as products of the surfactin production reactions. Sink-type boundary reactions were also added for every surfactin to remove them from the intracellular compartment [58]. To ensure surfactin production, lower bounds for each sink reaction were set at 0.0001 mmol gDW^−1^h^−1^. The FBA objective function was defined as the weighted average of the biomass reaction and the desired surfactin reaction in a 95:5 ratio. Details on medium components and their bounds can be found in Supplementary Table S2.

### 4.3 Modelling the media composition

The medium composition is defined by the molar concentration of each component. However, GEMs describe the medium with flux bounds in mmol gDW^−1^h^−1^. To convert between molar and flux units, we employed a scaling factor by matching glucose concentration and import fluxes to doubling times of *E. coli*. Specifically, a realistic doubling time for wild type *E. coli* in M9 is considered to be 42 min ± 12 min [59], which corresponds to a growth rate between 0.77 h^−1^ and 1.39 h^−1^. We therefore set the upper bound for the glucose flux to 10 mmol gDW^−1^h^−1^, as this is the value leading to a predicted growth rate of 0.85 h^−1^ in wild-type *E. coli*. To convert a glucose concentration of 20 mmol in M9 to 10 mmol gDW^−1^h^−1^, we required a conversion factor of 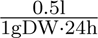, indicating that per gram of bacteria 500 ml of medium at the given concentration would be supplied over 24 hours. This conversion factor was applied to medium components in all case studies.

## Data and code availability

Python code for gsMOBO and data to reproduced the results can be found in Zenodo at https://doi.org/10.5281/zenodo.17798833.

## Acknowledgements

CM was supported by the United Kingdom Research and Innovation (grant EP/S02431X/1, UKRI Centre for Doctoral Training in Biomedical AI). CGC was supported by a PhD studentship from the Darwin Trust of Edinburgh.

## Supplementary Figures and Tables

**Table S1:**
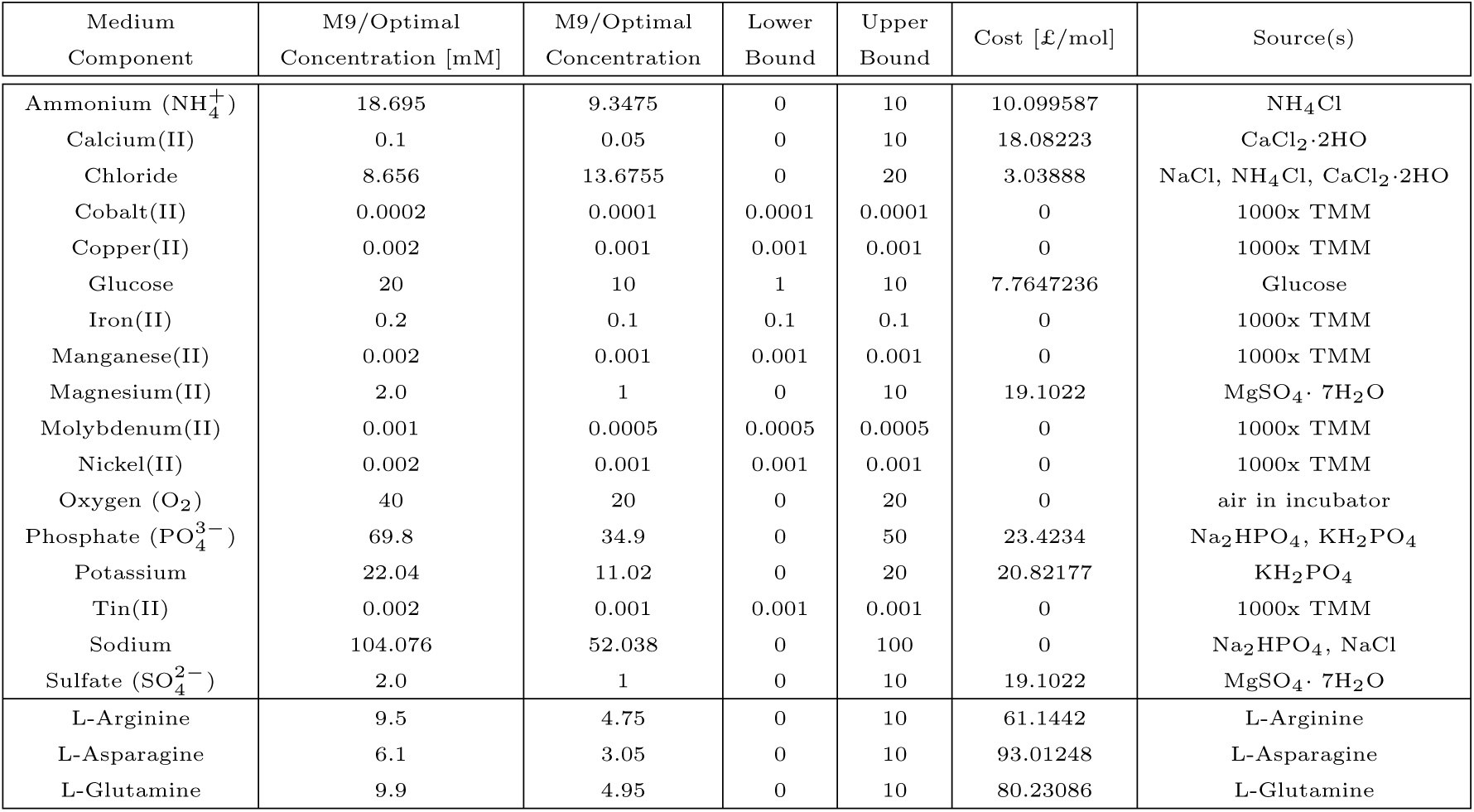
Medium components for case studies in *Escherichia coli*. Unless otherwise specified, concentrations and bounds are given in mmol gDW^−1^h^−1^. Prices for components with a fixed concentration (trace metals usually provided by a 1000x Trace Metals Mixture (1000x TMM)) and for components provided by the general infrastructure (carbon dioxide, water, hydrogen, oxygen) were set to zero. All other prices were taken from the Sigma Aldrich UK website in October 2024. The medium for iML1515 contains all listed components except for the amino acids. The medium for the iJO1366-antiEpEX-scFv model contains all other listed ingredients.

| Medium Component | M9/Optimal Concentration [mM] | M9/Optimal Concentration | Lower Bound | Upper Bound | Cost [£/mol] | Source(s) |
| --- | --- | --- | --- | --- | --- | --- |
| Ammonium (NH<sub>4</sub><sup>+</sup>) | 18.695 | 9.3475 | 0 | 10 | 10.099587 | NH<sub>4</sub>Cl |
| Calcium(II) | 0.1 | 0.05 | 0 | 10 | 18.08223 | CaCl<sub>2</sub>·2H<sub>2</sub>O |
| Chloride | 8.656 | 13.6755 | 0 | 20 | 3.03888 | NaCl, NH<sub>4</sub>Cl, CaCl<sub>2</sub>·2H<sub>2</sub>O |
| Cobalt(II) | 0.0002 | 0.0001 | 0.0001 | 0.0001 | 0 | 1000x TMM |
| Copper(II) | 0.002 | 0.001 | 0.001 | 0.001 | 0 | 1000x TMM |
| Glucose | 20 | 10 | 1 | 10 | 7.7647236 | Glucose |
| Iron(II) | 0.2 | 0.1 | 0.1 | 0.1 | 0 | 1000x TMM |
| Manganese(II) | 0.002 | 0.001 | 0.001 | 0.001 | 0 | 1000x TMM |
| Magnesium(II) | 2.0 | 1 | 0 | 10 | 19.1022 | MgSO<sub>4</sub>·7H<sub>2</sub>O |
| Molybdenum(II) | 0.001 | 0.0005 | 0.0005 | 0.0005 | 0 | 1000x TMM |
| Nickel(II) | 0.002 | 0.001 | 0.001 | 0.001 | 0 | 1000x TMM |
| Oxygen (O<sub>2</sub>) | 40 | 20 | 0 | 20 | 0 | air in incubator |
| Phosphate (PO<sub>4</sub><sup>3-</sup>) | 69.8 | 34.9 | 0 | 50 | 23.4234 | Na<sub>2</sub>HPO<sub>4</sub>, KH<sub>2</sub>PO<sub>4</sub> |
| Potassium | 22.04 | 11.02 | 0 | 20 | 20.82177 | KH<sub>2</sub>PO<sub>4</sub> |
| Tin(II) | 0.002 | 0.001 | 0.001 | 0.001 | 0 | 1000x TMM |
| Sodium | 104.076 | 52.038 | 0 | 100 | 0 | Na<sub>2</sub>HPO<sub>4</sub>, NaCl |
| Sulfate (SO<sub>4</sub><sup>2-</sup>) | 2.0 | 1 | 0 | 10 | 19.1022 | MgSO<sub>4</sub>·7H<sub>2</sub>O |
| L-Arginine | 9.5 | 4.75 | 0 | 10 | 61.1442 | L-Arginine |
| L-Asparagine | 6.1 | 3.05 | 0 | 10 | 93.01248 | L-Asparagine |
| L-Glutamine | 9.9 | 4.95 | 0 | 10 | 80.23086 | L-Glutamine |

**Table S2:** Medium components for case study in *Bacillus subtilis*. Unless otherwise specified, concentrations and bounds are given in mmol gDW^−1^h^−1^. The medium for iBsu1103 contains all the components from the M9 medium except from chloride and iron as only trace metal.

| Medium Component | M9/Optimal Concentration [mM] | M9/Optimal Concentration | Lower Bound | Upper Bound | Source(s) |
| --- | --- | --- | --- | --- | --- |
| Ammonium (NH<sub>4</sub><sup>+</sup>) | 18.695 | 9.3475 | 0 | 10 | NH<sub>4</sub>Cl |
| Calcium(II) | 0.1 | 0.05 | 0 | 10 | CaCl<sub>2</sub>·2H<sub>2</sub>O |
| Glucose | 20 | 10 | 1 | 10 | Glucose |
| Iron(II) | 0.2 | 0.1 | 0 | 10 | FeSO<sub>4</sub> |
| Magnesium(II) | 2.0 | 1 | 0 | 10 | MgSO<sub>4</sub>· 7H<sub>2</sub>O |
| Oxygen (O<sub>2</sub>) | 40 | 20 | 0 | 20 | air in incubator |
| Phosphate (PO<sub>4</sub><sup>3-</sup>) | 69.8 | 34.9 | 0 | 50 | Na<sub>2</sub>HPO<sub>4</sub>, KH<sub>2</sub>PO<sub>4</sub> |
| Potassium | 22.04 | 11.02 | 0 | 20 | KH<sub>2</sub>PO<sub>4</sub> |
| Sulfate (SO<sub>4</sub><sup>2-</sup>) | 2.0 | 1 | 0 | 10 | MgSO<sub>4</sub>· 7H<sub>2</sub>O |
| Carbon Dioxide (CO<sub>2</sub>) | N/A | 0 | 0 | 10 | produced in fermentation |
| Protons (H<sup>+</sup>) | N/A | 0 | 0 | 10 | produced in fermentation |
| Water (H<sub>2</sub>O) | N/A | N/A | 0 | 100 | solvent |

**Figure S1:**
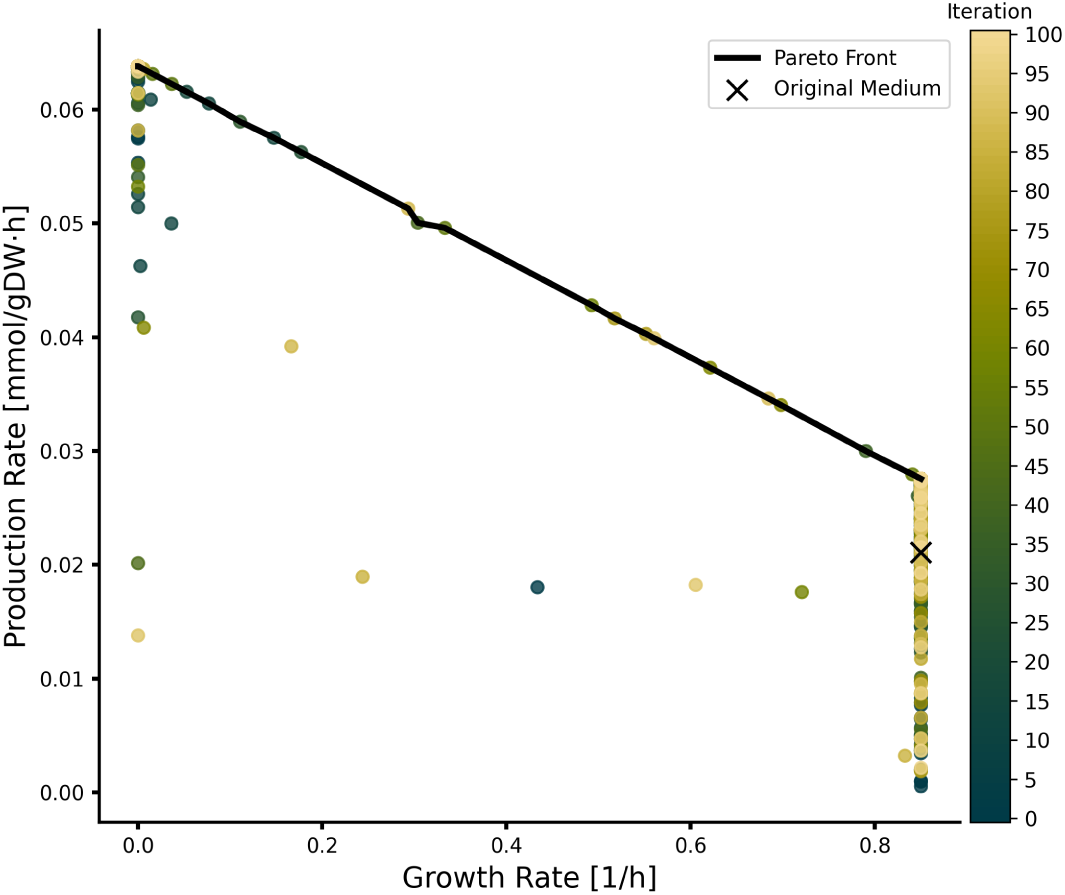
Double-objective optimization of antibody-producing *E. coli* GEM medium conditions. The optimization, with the objective to maximize both growth rate and production rate, was run for N=100 iterations, a batch size of 15, and 50 random starting medium compositions.

**Figure S2:**
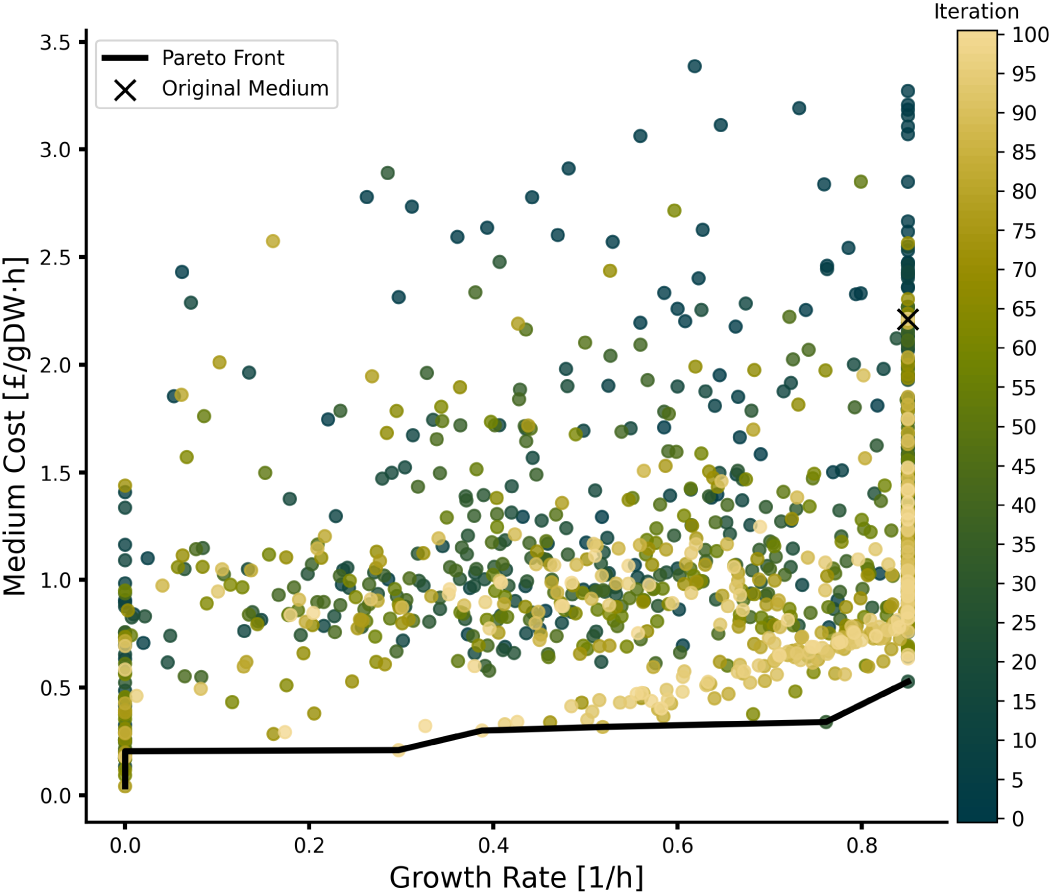
Double-objective optimization of antibody-producing *E. coli* GEM medium conditions. The optimization, with the objective to maximize the growth rate while minimizing the cost, was run for N=100 iterations, a batch size of 15, and 50 random starting medium compositions.

**Figure S3:**
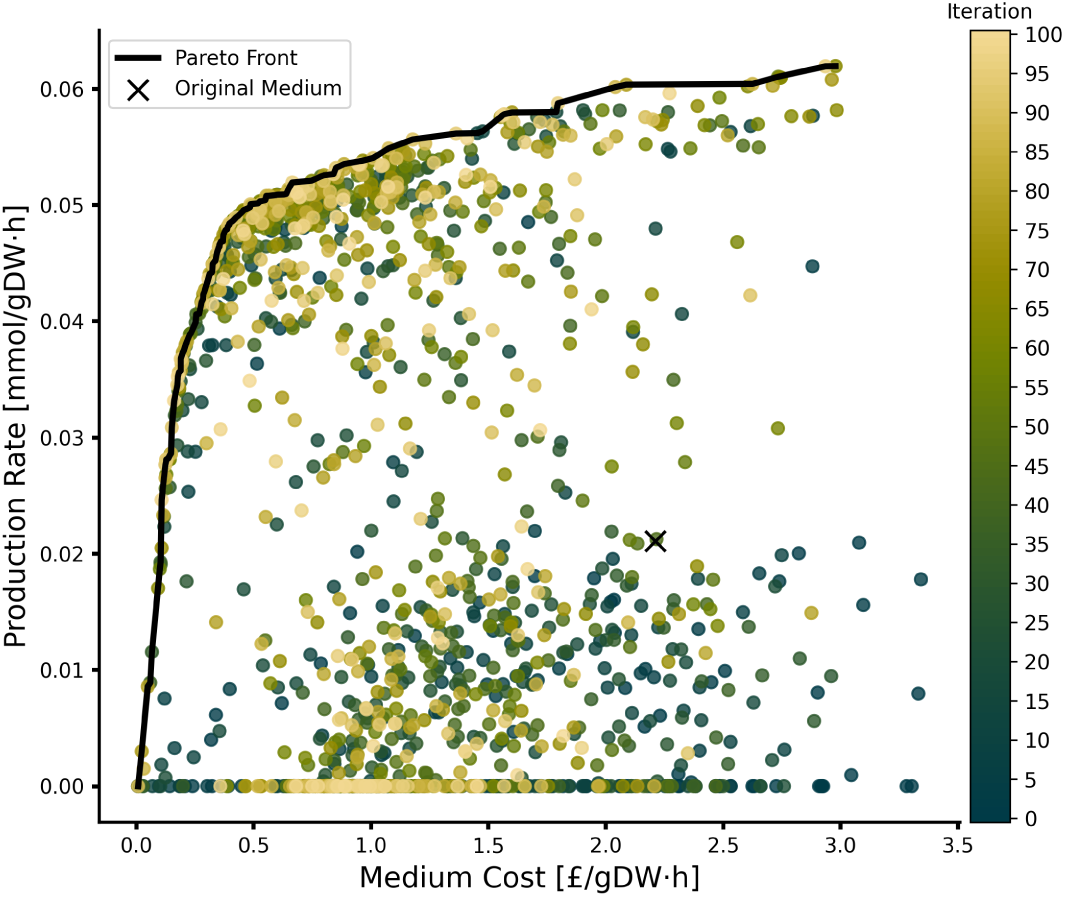
Double-objective optimization of antibody-producing *E. coli* GEM medium conditions. The optimization, with the objective to maximize the production rate while minimizing the cost, was run for N=100 iterations, a batch size of 15, and 50 random starting medium compositions.

**Figure S4:**
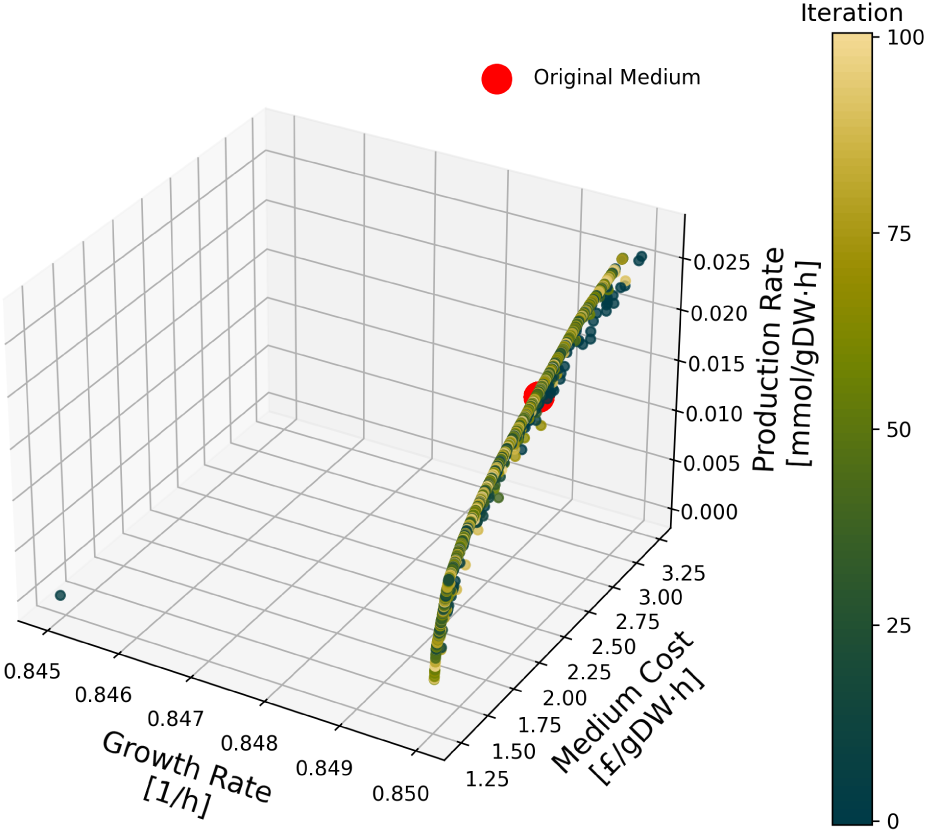
Triple-objective optimization of antibody production in *E. coli*. This optimization is analogous to Figure 3, but with fixed M9 medium and only the amino acid components as decision variables.

